# De novo transcriptome sequencing of *Serangium japonicum* (Coleoptera: Coccinellidae) and application of two assembled unigenes

**DOI:** 10.1101/605410

**Authors:** Ya Hui Hu, Yong Liu, Lin Wei, Hao Tao Chen

## Abstract

The ladybird beetle *Serangium japonicum* is an important predator of whiteflies. Although some ecological and biological characteristics of this predator have been studied, the only molecular data for the ladybird beetle at the NCBI website has been submitted by us. To yield gene sequences of the ladybird beetle, paired-end sequencing was used to sequence its transcriptome. Real-time PCR was used to validate differences in the quantity of RNA expressed by the *Krueppel homolog 1* (*Kr-h1*) gene in summer versus winter samples. To find the effective molecular barcode, the annotated c*ytochrome c oxidase subunit I* (*COX1*) gene fragments were amplified from several ladybird beetle populations. Analysis yielded 191,246 assembled unigenes, of which 127,016 (66.4%) were annotated. The differential expression of *Kr-h1* in summer versus winter suggests that *S. japonicum* can successfully overwinter because the adults enter diapause. The annotated *COX1* gene can be used to distinguish *S. japonicum* from other ladybird species. These gene sequences are currently available in the National Center for Biotechnology Information (NCBI), and will facilitate the study of molecular mechanism in *S. japonicum*.

## Introduction

Whiteflies are notable pests that prey on many horticultural crops (Ren et al., 2001; Ren et al., 2011). *Serangium japonicum* have been reported as an effective or potential predator of several types of whiteflies, such as *Bemisia tabaci* (Ren et al., 2001; Sahar and Ren 2004; Li et al., 2015), *Dialeurodes citri* (Kaneko, 2017), and *Aleurocanthus camelliae* (Ozawa and Uchiyama 2016). *S. japonicum* has been studied for its biological characteristics (Yao et al., 2005; Fatiha et al.□2008; Yao et al., 2011; Li et al., 2014; Hu et al., 2016; Kaneko, 2017) and response to insecticides (Hu et al., 2009; He et al., 2012; Zhao et al., 2012; Yao et al., 2015) and juvenile hormone analog (Li et al., 2015). Gene sequences from *S. japonicum* have not been reported by others.

Next-generation RNA sequencing has been used to assess the molecular mechanisms underlying processes in insects (Zhang et al., 2014; Qi et al., 2016). Compared with other molecular technologies, next-generation RNA sequencing has a lower cost.

Transcriptome sequencing can effectively identify molecular markers (Parra-Gonzalez et al., 2012). The assembled genes from the transcriptome can be expressed differently in selected tissues or organs, and at various developmental stages (Fu et al., 2016). Moreover, when a reference genome is unavailable, next-generation RNA sequencing can effectively obtain the annotated gene sequences of a species (Martin and Wang 2011) via BLAST (basic local alignment search tool) with other species’ gene sequences in several public databases.

Diapause is a behavior that allows insects to adapt to an unfavorable environment. Cold dormancy temperatures (5–8□) lasting for 30 days can be fatal to *S. japonicum* adults if diapause is not induced before they are exposed to such temperatures (Hu et al., 2016). Insects in diapause can survive in cold weather for several months, particularly if they have accumulated fatty material in the body before overwintering (Hahn and Denlinger 2011). Juvenile hormone (JH) can control developmental transitions in insects, including diapause. The *Krueppel homolog 1* (*Kr-h1*) gene plays an important role in diapause, inducing transcription of *JH* (Jindra et al., 2013; Jindra et al., 2015). This project aimed to explore whether there is a difference of *Kr-h1* expression in *S. japonicum* in summer versus winter.

The *cytochrome c oxidase subunit I* (*COX1*) gene can be used to identify different biological species (Hebert et al. 2004). In insects, COX1 has been successfully used to distinguish between different whitefly species or biotypes and different geographical populations within a whitefly species (Ren et al., 2011). *S. japonicum* are difficult to distinguish from *Delphastus catalinae* because they have similar morphological features. Additionally, *S. japonicum* is distributed throughout many provinces in China and several other countries. It is expected that a COX1 sequence fragment can be regarded as an effective DNA barcode of *S. japonicum*.

## Materials and methods

### RNA extraction and sequencing

*S. japonicum* samples were collected from eggplants at the Academy of Hunan Agricultural Science, Changsha, China 3 times during winter 2016 and 3 times during summer 2017. Each time, 7 *S. japonicum* adults were collected from plants as a biological sample, for a total of 42 adults. Total RNA was extracted from *S. japonicum* with the EasyPure RNA kit (Transgen Biotech, Beijing, China). RNA was sequenced by Sangon Biotech (Shanghai) Co., Ltd., Shanghai, China, on an Illumina HiSeq 2500.

### De novo transcriptome assembly

Raw reads from 6 biological samples were cleaned by removing adapter sequences (forward: AGATCGGAAGAGCACACGTCTGAAC; reverse: AGATCGGAAGAGCGTCGTGTAGGGA) and both sides bases Q<20 from reads, removing reads with unknown nucleotides “N”, and removing reads <35 nucleotides in length. The clean reads from both groups were assembled *de novo* using Trinity (Haas et al. 2013). The Trinity default parameter setting was used, except for min_kmer_cov. Trinity treated the cleaned reads via 3 steps: Inchworm, Chrysalis, and Butterfly. None of assembled sequences <200 nucleotides were regarded as unigenes.

### Functional annotation

The generated unigenes were annotated based on the following 5 databases: the National Center for Biotechnology Information (NCBI) non-redundant protein database (Nr), the NCBI non-redundant nucleotide database (Nt), Swiss-Prot, the Eukaryotic Orthologous Groups (KOG) database, and the Kyoto encyclopedia of genes and genomes (KEGG), with E-value <1 × 10^-5^. The best-aligned results were used to decide the sequence direction and coding sequence of unigenes. If results of different databases conflicted with each other, the following order of priority was employed: Nr, Nt, Swiss-Prot, KEGG, and COG. Even if unigenes were not annotated in any of listed databases, their sequence direction and coding sequence would be predicted by TransDecoder (v3.0.1) (http://transdecoder.github.io/). Distribution of similar species was analysed based on Nr database annotation (Shi et al., 2011).

### Kr-h1 expression in summer and winter

The level of unigene expression was estimated by measuring transcripts per million reads (TPM) (Patro et al. 2017). In addition, the DESeq test was used to identify differentially expressed genes between the respective TPMs of summer versus winter samples, with p≤0.05 and ≥2-fold change. One microgram of RNA was employed for first-strand cDNA synthesis with a RevertAid Premium Reverse Transcriptase kit (Thermo Scientific™) used according to the manufacturer’s instructions. Real-time polymerase chain reaction (PCR) was performed with SG Fast qPCR Master Mix (High Rox) at 95°C for 3 min, followed by 45 cycles of 95°C for 7 s and 56°C for 10 s. The primers used were 5’-3’sequence CTTCATCTTGCTGGAATCTCC and 5’-3’ sequence AATAGCTCCTGCTAATACTGGTAA. Melting curves were analysed from 60°C to 95°C to detect nonspecific product amplification. The assembled gene_id: N79847_c1_g1 of *S. japonicum* was used as an internal control. Data analysis was carried out by the 2^-ΔΔCT^ method.

### COX1 gene as barcode investigation of the geographic populations

The investigated populations of *S. japonicum* were distributed in Changsha in Hunan province, Mianyang in Sichuan province, and Nanjing in Jiangsu province. According to the transcriptome sequencing, assembled unigenes, and annotation from Nr, we designed the following primer pair: 5’-3’: tattttctttttggactttg, 5’-3’: gtaatgttgctaatcaagaaaa. These primers amplified a 980-nucleotide COX1 gene fragment from 3 populations via PCR. Sequences from each of the 3 *S. japonicum* populations were aligned.

### Accession numbers

The raw reads produced in this study can be obtained in NCBI by searching the project numbers PRJNA376265 and PRJNA430037; BioSample numbers SAMN06347100, SAMN08365344, and SAMN08365343; or with the accession codes SRR5277648, SRR6473305, SRR6473306, SRR6473307, SRR6473308, and SRR6473309. The assembled unigene sequences have been submitted to the Transcriptome Shotgun Assembly sequence database with the accession code GGMU00000000.

## Results

### Sequencing and assembly

More than 142 million raw reads were obtained from each group of samples (Table 1), resulting in 133.29 million cleaned reads (92.4% of the raw reads) for the 3 summer adult samples, and 140.34 million (93.7% of the raw reads) for the 3 winter adult samples. The additional file 1 showed raw and cleaned reads satisfied from individual biological sample. Analysis yielded 191,246 unigenes with an average length of 524 nucleotides and an N50 of 681 nucleotides; 54,929 (28.7%) and 21,351 (11.2%) unigenes were longer than 500 nt and 1000 nt, respectively (Figure 1).

**Table 1.**
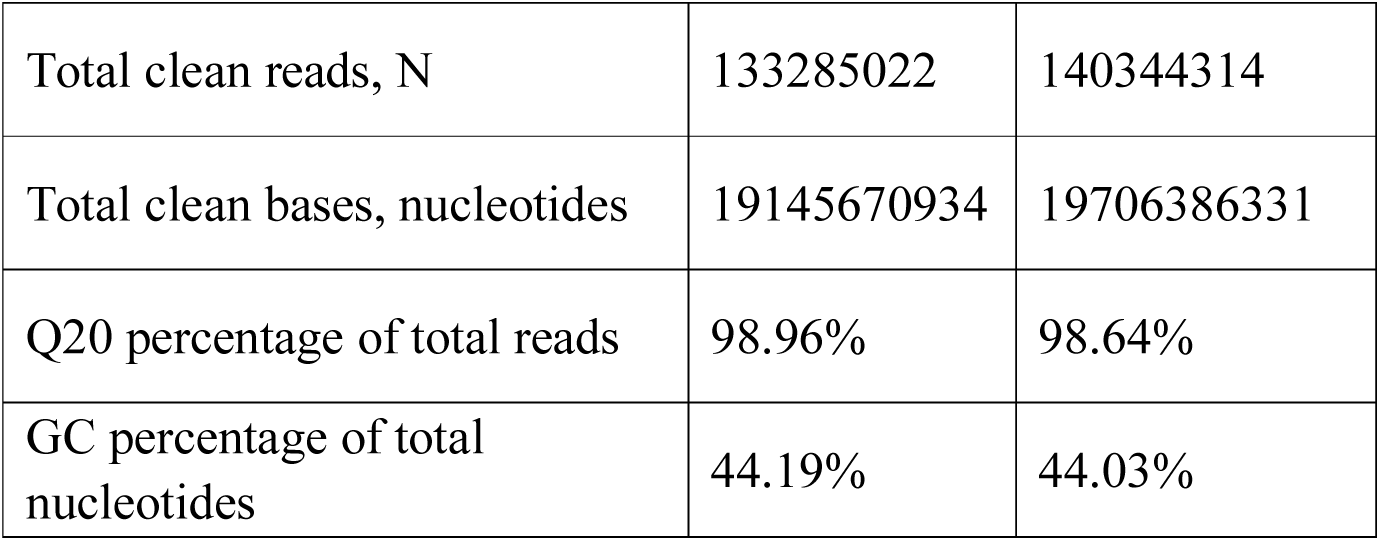
Sequencing data from *S. japonica* samples collected in summer versus winter

**Figure 1.**
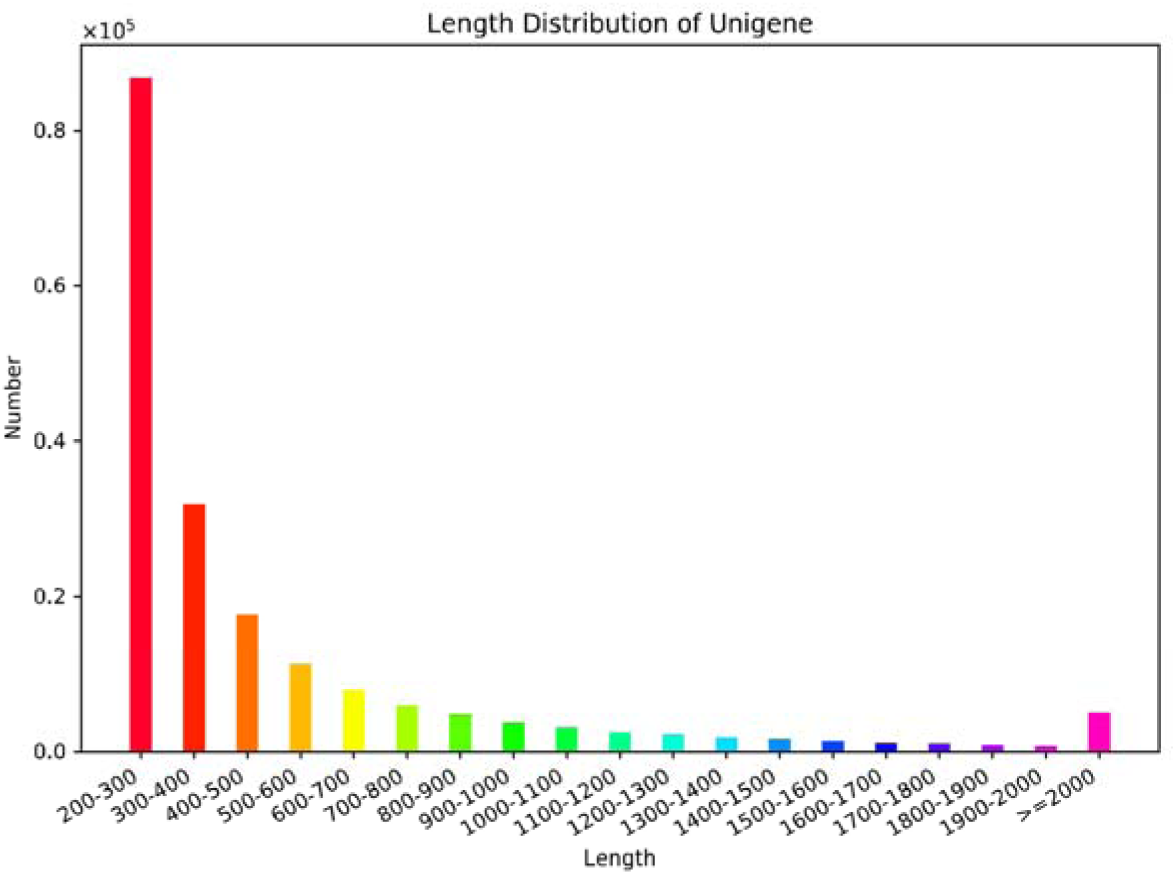
Length distribution of the obtained unigenes

### Functional annotation

Unigenes were annotated according to the Nr, Nt, Swiss-Prot, COG, KEGG database. After searches in these 5 databases, a total of 127,016 (66.4% of 191,246) unigenes could be annotated: 89,349 in Nr, 85,085 in Swiss-Prot, 79,088 in Nt, 62,877 in the KOG database, and 10,554 in KEGG.

According to the Nr database annotation, genes from *S. japonicum* matched *Tribolium castaneum* most similarly (Figure 2). According to search results from the KOG database, the 3 largest categories were general function prediction (8503; 13.52%); signal transduction mechanisms (8175; 13.00%); and posttranslational modification protein turnover chaperones (6548; 10.41%) (Figure 3). According to the KEGG, a total of 24,653 unigenes were identified by 309 KEGG pathways; the most represented were ribosomal pathways (847 unigenes, 3.4%), followed by oxidative phosphorylation (502; 2.0%), carbon metabolism (431; 1.7%), and biosynthesis of amino acids (414; 1.7%). These functional annotations of unigenes provide a basis for studying the molecular mechanisms underlying the biological characteristics of *S. japonicum*.

**Figure 2.**
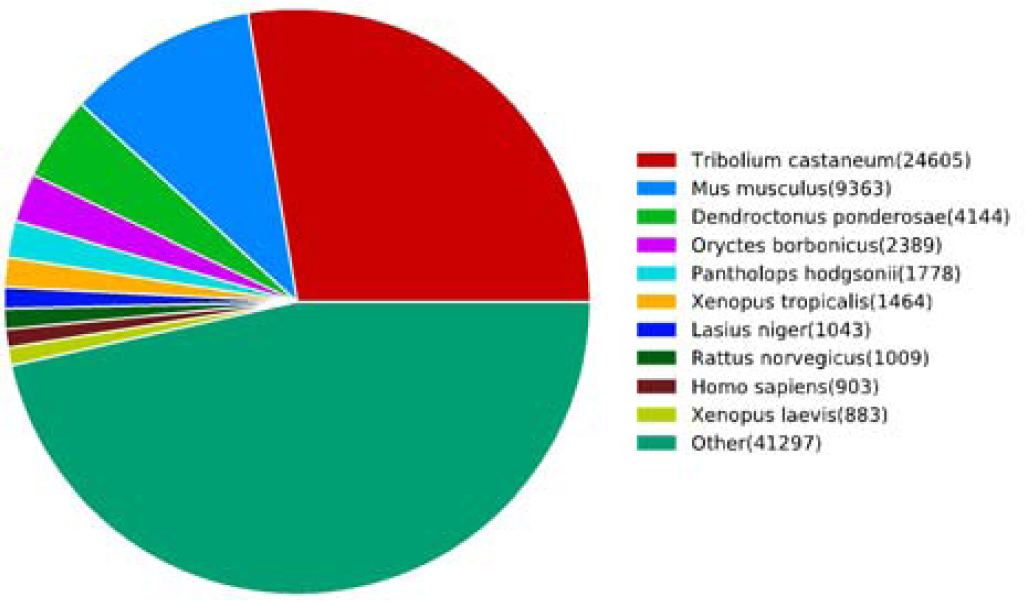
Distribution of species in the NCBI non-redundant protein database

**Figure 3.**
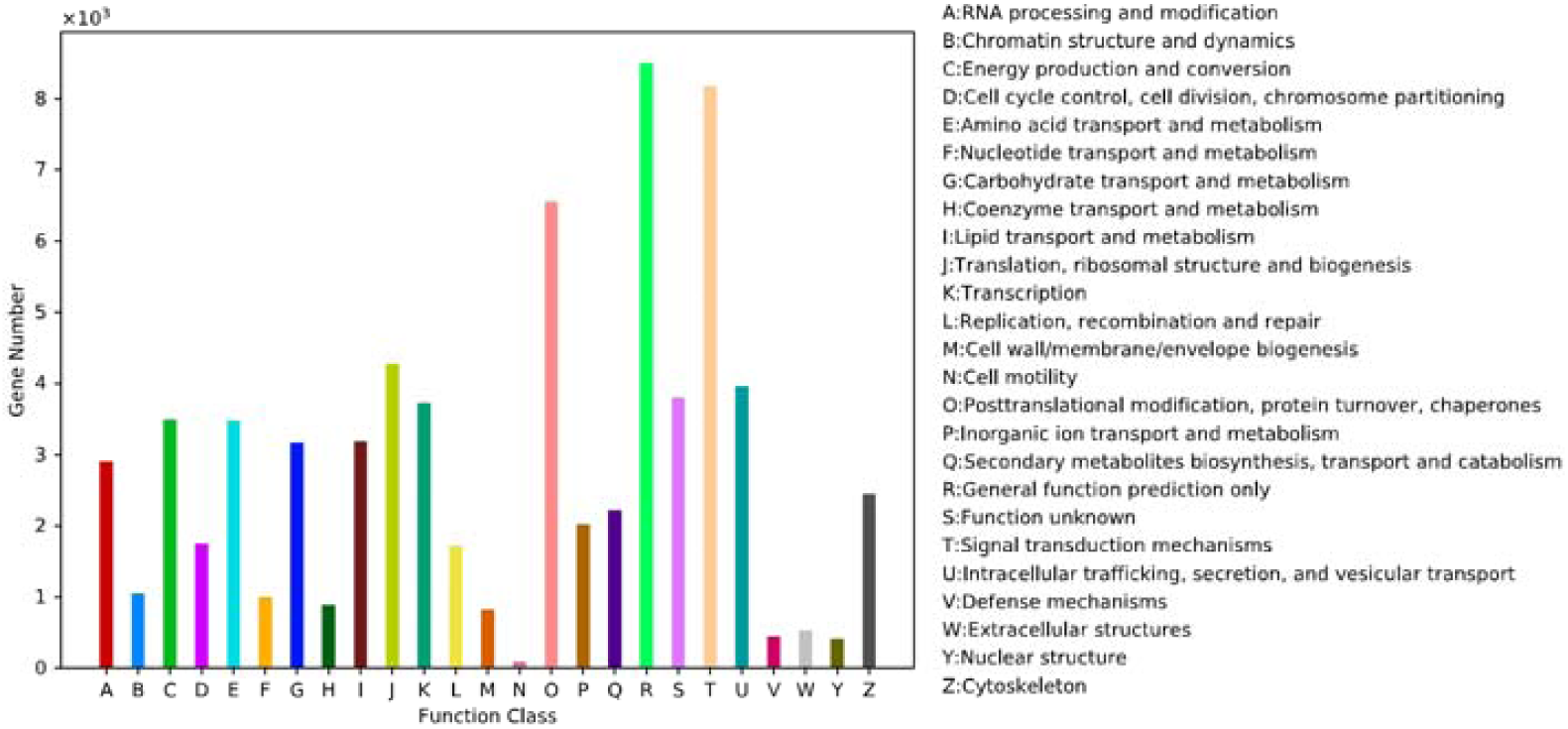
Eukaryotic orthologous group (KOG) categories

### Kr-h1 expression in summer and winter

Expression of the *Kr-h1* gene (Gene id: TRINITY_DN74054_c1_g1) was down-regulated in *S. japonicum* in winter compared with summer (mean TPM values: 2.5 in winter vs. 33.1 in summer; log2Foldchange 3.75; p=0.038). The q-PCR results for the *Kr-h1* gene showed log2Foldchange=3.02 (p=0.021). The additional file 2 recorded other differentially expressed genes according to high-throughput sequencing.

### Application of the COX1 gene

The assembled sequence (Gene id: TRINITY_DN81324_c0_g4) was annotated as the gene for *COX1* according to the Nr, Nt, Swiss-Prot, and KOG databases. The PCR results showed the sequences were identical in *S. japonicum* in Changsha, Mianyang, and Nanjing according to a 928-nucleotide COX1 fragment. In the NCBI nucleotide database, the assembled sequence had the highest identity value (91%) with the sequence ID KP829591.1 from *Serangium sp.* (Table 2).

**Table 2.** Taxonomy BLAST reports from the National Center for Biotechnology Information (NCBI) database

| Organism | BLAST Name | Score | Number of Hits |
| --- | --- | --- | --- |
| Holometabola | insects |  | 104 |
| Coleoptera | beetles |  | 90 |
| Polyphaga | beetles |  | 87 |
| Cucujiformia | beetles |  | 57 |
| Cucujoidae | beetles |  | 21 |
| Coccinellidae | beetles |  | 10 |
| Serangium sp. CO631 | beetles | 1628 | 1 |
| Scymninae sp. 2ACP-2013 | beetles | 1567 | 1 |
| Propylea japonica | beetles | 1506 | 1 |

RID: 4TZB21ZH015 (Expires on 01-28 07:37 am)
Query ID: lcl|Query\_11215
Molecule type: nucleic acid
QueryLength: 1643
Database Name: nr
Description: Nucleotide collection (nt)
Program: BLASTN 2.8.1+ Citation

## Discussion

### Kr-h1, diapause, and overwintering

Temperatures of 5–8□ for 30 days can be fatal to *S. japonicum* adults if they do not enter diapause before exposure to the cold; in contrast, adults in diapause can survive for several months (Hu et al., 2016). JH acts together with the Methoprene-tolerant and Germ cell-expressed bHLH-PAS transcription factors (which act as potential JH receptors) to directly induce *Kr-h1* expression (Minakuchi et al., 2008; Lozano and Belles 2011; Kayukawa et al., 2012). Levels of the JH esterase and JH were low in diapause adults compared with non-diapause adults (Qi et al., 2016). The down-regulation of *Kr-h1* expression suggested that *S. japonicum* adults in winter were in diapause, with low JH levels. After diapause, the insect can successfully overwinter in low-temperature conditions because of the accumulation of fatty acids, trehalose, and other energy sources (Hahn and Denlinger 2011; Tang et al., 2017).

The *Kr-h1* gene negatively regulates ecdysone biosynthesis by directly inhibiting the transcription of steroidogenic enzymes (Liu et al., 2018; Zhang et al., 2018). A hormone receptor also acts as a repressor of ecdysone biosynthesis in *Drosophila melanogaster* (King-Jones et al., 2005; Qu et al., 2011). Other hormone receptors may inhibit ecdysone biosynthesis might be inhibited in *S. japonicum* adults in diapause (i.e., with low *Kr-h1* expression). Future studies will explore whether ecdysone biosynthesis is inhibited during pre-diapause by high *Kr-h1* expression in *S. japonicum*. The relationship between *Kr-h1* and other hormone receptors in insects is also of increasing interest.

### S. japonicum species and population

*Serangium sp., Scymninae sp.*, and *Propylea japonica* have high *COX1* gene identity with *S. japonicum*. The COX1 from *S. japonicum* was morphologically consistent with species taxonomy. It was easy to accurately distinguish *S. japonicum* from the morphologically similar species *Delphastus catalinae* by aligning *COX1* genes from the 2 species. In contrast to its prey, the whitefly (Ren et al., 2011□Kanmiya et al., 2011), *S. japonicum* has only one population throughout several provinces in China, based on examination of the *COX1* gene from their samples. We suggest *S. japonicum* can be widely applied to control whitefly populations in many regions after the predator population is expanded/propagated.

## Conclusions

*S. japonicum* is an effective predator of whiteflies. However, better use of this species requires thorough study of the molecular mechanisms underlying diapause, overwintering, and other biological characteristics. The study of molecular mechanisms of this predatory beetle is hindered by the scarcity of gene sequence data. The Illumina Hiseq2500 sequencing platform was used to sequence the *S. japonicum* transcriptome, yielding 191,246 assembled unigenes, of which 127,016 (66.4%) were annotated. This study identified an abundance of genes in *S. japonicum*. Annotation of unigenes would facilitate understanding of the mechanisms underlying biological characteristics in this species. The differential expression of *Kr-h1* in Summer/Winter suggests that *S. japonicum* can successfully overwinter because the adults enter diapause. The annotated *COX1* gene can be used to distinguish *S. japonicum* from other ladybird species. We were delighted to learn that a single *S. japonicum* population can be used in multiple regions.

## Acknowledgments

This work was supported by the National key R&D Program of China (2017YFD0201000), the Hunan Talent Project (2016 RS 2019), and the Hunan Vegetable Industry Technology System (2015-2019).

